# A thermophilic phage uses a small terminase protein with a fixed helix-turn-helix geometry

**DOI:** 10.1101/851345

**Authors:** Janelle A. Hayes, Brendan J. Hilbert, Christl Gaubitz, Nicholas P. Stone, Brian A. Kelch

## Abstract

Tailed bacteriophage use a DNA packaging motor to encapsulate their genome during viral particle assembly. The small terminase (TerS) component acts as a molecular matchmaker by recognizing the viral genome as well as the main motor component, the large terminase (TerL). How TerS binds DNA and the TerL protein remains unclear. Here, we identify the TerS protein of the thermophilic bacteriophage P74-26. TerS^P76-26^ oligomerizes into a nonamer that binds DNA, stimulates TerL ATPase activity, and inhibits TerL nuclease activity. Our cryo-EM structure shows that TerS^P76-26^ forms a ring with a wide central pore and radially arrayed helix-turn-helix (HTH) domains. These HTH domains, which are thought to bind DNA by wrapping the helix around the ring, are rigidly held in an orientation distinct from that seen in other TerS proteins. This rigid arrangement of the putative DNA binding domain imposes strong constraints on how TerS^P76-26^ can bind DNA. Finally, the TerS^P76-26^ structure lacks the conserved C-terminal β-barrel domain used by other TerS proteins for binding TerL, suggesting that a well-ordered C-terminal β-barrel domain is not necessary for TerS to carry out its function as a matchmaker.

## INTRODUCTION

Viruses infecting all domains of life, from bacteria to eukaryotes, replicate and encapsulate their genetic material to create infectious particles. For viruses with large genomes, transporting genetic material into the capsid is an energetic challenge, and many viruses have evolved motor systems to accomplish this task. Viruses with concatemeric double-stranded DNA genomes, such as herpesviruses, and most phage use a motor known as a ‘terminase motor’. Terminase motors are composed of three components: a ‘portal’ channel, a ‘small terminase’ DNA recognition protein, and a ‘large terminase’ that contains both nuclease and ATPase activities (Feiss and Rao, 2011). The portal, which is embedded within the capsid wall, acts as an adaptor to connect the capsid to the large terminase. The large terminase (TerL) binds portal and pumps DNA through its pore into the capsid. In order for this packaging step to occur, the motor must first specifically recognize the viral genome. This DNA-recognition task is performed by the small terminase (TerS), which binds a recognition sequence known as ‘cos’ or ‘pac’ and transfers the DNA to TerL for subsequent cleavage and packaging. Cos- and pac-containing phage are distinct in their cleavage mechanisms, as cos-phage only cleave at the cos site between genomes, whereas pac-containing phage solely use the pac site for packaging initiation, with the position of subsequent cleavage events dependent on a head-full sensing mechanism. It has been demonstrated that TerS has an important role in packaging initiation, as aberrant pac recognition impedes faithful genome packaging (Casjens et al., 1992; Schmieger, 1972).

Despite several decades of investigation, how TerS binds to pac is still unclear. In many viral genomes, the pac site is located within the gene for TerS itself (Baumann and Black, 2003; Casjens et al., 1987; Chai et al., 1995; Leavitt et al., 2013; Roy et al., 2012; Wu et al., 2002). The pac site of phage SPP1 appears to be flexible, suggesting a role for DNA bending in TerS recognition (Chai et al., 1995). Further clues for the DNA binding mechanism come from structures of TerS proteins. All currently known pac-recognizing TerS proteins multimerize into a ring with a central pore (Buttner et al., 2012; Roy et al., 2012; Sun et al., 2012; Zhao et al., 2010). In some of these assemblies, such as *Shigella flexneri* phage Sf6 and *Bacillus subtilis* phage SF6, the pore is too narrow to accommodate double-stranded DNA binding (Suppl. Table 1) (Buttner et al., 2012; Zhao et al., 2010). In these structures, the outward-facing N-terminal domain is a helix-turn-helix motif, a common DNA-binding domain. Studies of Sf6 TerS indicate that mutation of this region of the protein abrogates DNA binding, suggesting a nucleosome-like wrapping mechanism (Zhao et al., 2012). The exception to this model is the TerS structure of phage P22. In P22, the perimeter of the ring lacks the helix-turn-helix motif, and the pore is wide enough to accommodate DNA (Roy et al., 2012). This finding led to a second ‘threading’ model in which DNA binds in the center of the ring, traversing through the pore (Suppl. Table 1).

Regardless of the location of the DNA binding regions, all known TerS rings retain the same mushroom-like shape with a C-terminal β-barrel. TerS interacts with TerL using this β-barrel region, which is conserved in all TerS structures to date (Gao and Rao, 2011; Roy et al., 2012). TerS binding increases TerL’s ATPase activity while inhibiting nuclease activity (Baumann and Black, 2003; Gual et al., 2000; Leffers and Rao, 2000; Roy et al., 2012; Sun et al., 2012), suggesting that TerS has a regulatory effect on DNA packaging. Additionally, the β-barrel can control TerS assembly, as removing it causes polydisperse ring formation (Buttner et al., 2012; Sun et al., 2012). Therefore, the C-terminal β-barrel has been hypothesized to be important for both TerS oligomerization and regulation of TerL activity.

In past studies, we have used the thermophilic phage model system P74-26 to probe the mechanisms behind different stages of the viral life cycle (Hilbert et al., 2015, 2017; Stone et al., 2018). Here, we identify and characterize the small terminase gene of phage P74-26, hereafter known as TerS^P74-26^. TerS^P74-26^ binds DNA and both activates ATPase and inhibits nuclease activity of TerL^P74-26^. We report symmetric and asymmetric cryo-EM reconstructions of TerS^P74-26^ to overall resolutions of 3.8 Å and 4.8 Å resolution, respectively. Our structures show that TerS^P74-26^ retains the N-terminal helix-turn-helix motif, while also having a wide enough pore for DNA binding. In comparison to other TerS proteins, the helix-turn-helix domain is in a distinct conformation, with implications for the DNA binding mechanism. Finally, the C-terminal region of P74-26 TerS is unstructured, indicating that the β-barrel fold is not strictly conserved, nor is it essential for regulating P74-26 TerL activity.

## RESULTS

### Identification of P74-26 gp83 as the small terminase (TerS)

To investigate how thermophilic small terminase proteins recognize the viral genome, we sought to identify and characterize the TerS of P74-26 phage. TerS proteins commonly exhibit low sequence conservation, which can make their identification challenging. However, synteny can be used to identify the gene, as the small terminase gene often directly precedes the large terminase gene. Because gene 84 encodes the large terminase (Minakhin et al., 2008), we hypothesized that the gp83 protein is TerS. Although gp83 has low sequence homology to any known TerS protein (closest relative being T4 TerS, which retains 19% identity), its length of 171 amino acids is similar to that of known TerS proteins.

To further verify its identity, the putative TerS protein was recombinantly expressed and purified to homogeneity (Figure 1A). Size-exclusion multi-angle light scattering (SEC-MALS) shows gp83 assembles into a stable 9-mer complex, with a measured molecular mass of 170 kDa (compared to 171 kDa calculated by sequence) and a polydispersity index of 1.000, indicating a monodisperse assembly (Figure 1B). The oligomerization state of gp83 is consistent with that of mesophilic TerS proteins, which assemble into 8 to 11 subunit oligomers (Buttner et al., 2012; Roy et al., 2012; Sun et al., 2012; Zhao et al., 2010).

**Figure 1.**
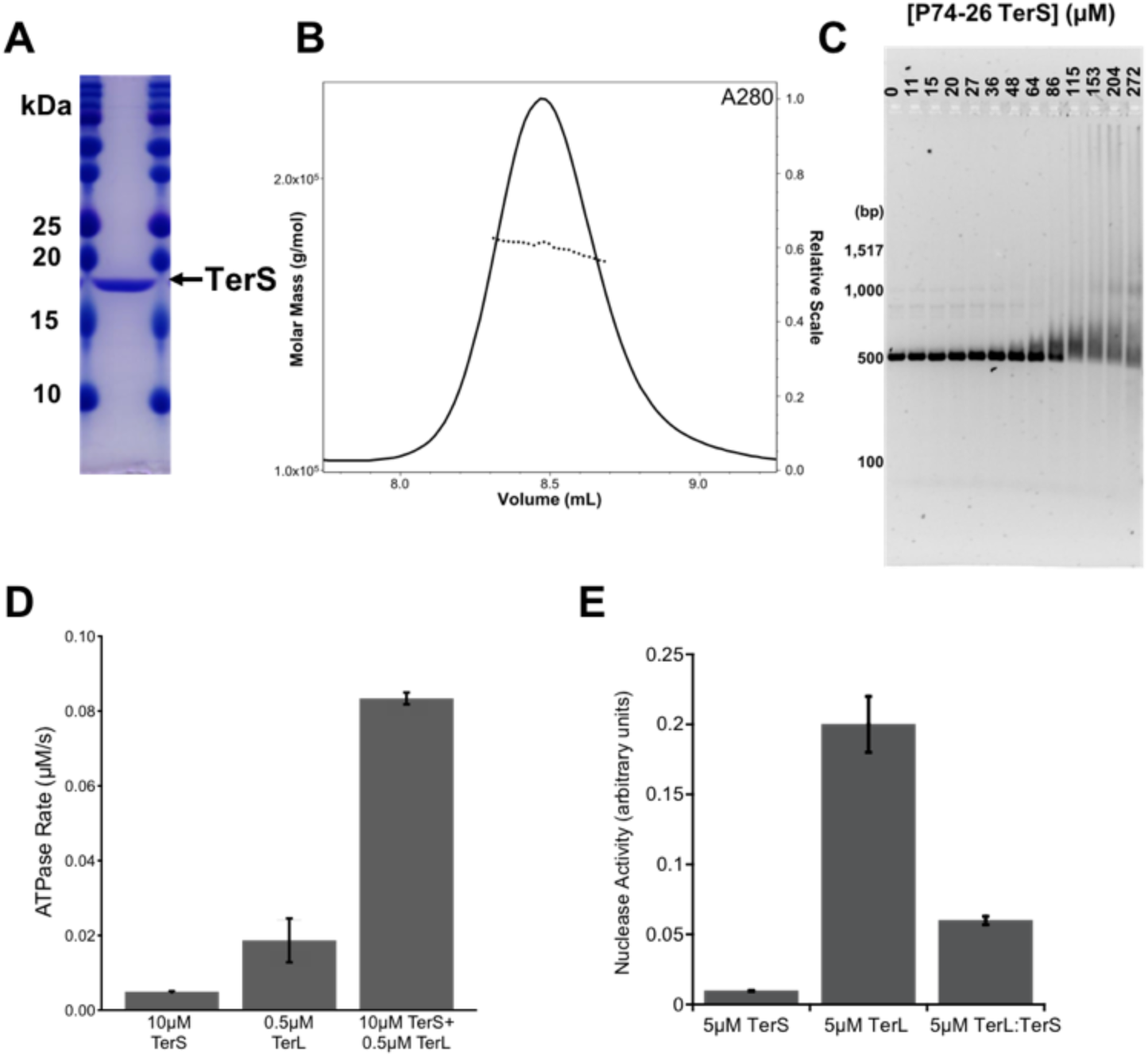
Characterization of TerS gp83. **(A)** SDS-PAGE gel of purified P74-26 gp83. **(B)** SEC-MALS of P74-26 gp83. The UV absorbance at 280 nm wavelength is shown. The measured molecular mass of the complex is 170 kDa, compared to 171 kDa calculated from sequence of a 9-mer. The polydispersity index is 1.000. **(C)** P74-26 gp83 binds DNA with weak affinity. Titrating P74-26 gp83 from 0 to 272 µM (monomer) with 50 ng of the P74-26 gp83 gene shows TerS has a low affinity for DNA. **(D)** P74-26 gp83 increases ATPase activity of TerL^P74-26^ 4.4-fold. **(E)** P74-26 gp83 decreases TerL^P74-26^ nuclease activity 3.3-fold.

To determine if gp83 binds DNA like other TerS proteins, we performed electromobility shift assays. Because many other TerS oligomers recognize a sequence within their own gene (Baumann and Black, 2003; Casjens et al., 1987; Chai et al., 1995; Leavitt et al., 2013; Roy et al., 2012; Wu et al., 2002), we used the P74-26 gp83 DNA sequence to evaluate DNA binding. The gp83 complex binds DNA weakly, as indicated by smearing within the gel (Figure 1C). Low DNA binding affinity is commonly seen in other TerS proteins (Greive et al., 2016; Zhao et al., 2012).

We also find that gp83 modulates the enzymatic activities of TerL. Upon mixing gp83 with TerL^P74-26^, ATPase activity increases 4.4-fold (Figure 1D). This suggests a direct interaction between TerL and gp83, as no DNA is present in the experiment. gp83 also inhibits TerL nuclease activity 3.3-fold (Figure 1E). The modulation of TerL enzymatic activities is consistent with previous studies of TerS proteins from other phages (Alam et al., 2008; Gual et al., 2000; Leffers and Rao, 2000; Sun et al., 2012). Taken together, our results identify gp83 as the TerS of P74-26.

### The structure of TerS^P74-26^

We next used electron microscopy (EM) to determine the structure of TerS^P74-26^. Negative stain EM shows homogenous TerS particles with even distributions of top and side views (Suppl. Figure 1A). From 2D classification, we observe that TerS^P74-26^ forms a ring-shaped assembly with a central pore. To further elucidate the structure of TerS^P74-26^, we prepared samples of the complex for single-particle reconstruction by cryo-EM. Unlike negative stain samples, cryo-EM samples show strong preferred orientation for the top and bottom views of the ring and slight aggregation (Suppl. Figure 1B). The lack of side views severely hampers initial structure determination, and the middle portion of the ring cannot be resolved (Suppl. Figure 1C).

To increase particle side views, we used a combination of sample additives and tilted data collection. Out of the numerous additives tested, amphipol A8-35 had the greatest effect on particle view distribution. After collecting a set of un-tilted images, we used a 30° tilt to obtain additional particle views (Suppl. Figures 2A-C). Initial 3D classification of the combined datasets produces six different classes, several of which are of particular interest (Figure 2A). Classes 1 and 2, which account for over 50% of all particles, show apparent 9-fold symmetry. Asymmetric refinement of these combined classes generates a reconstruction with an overall resolution of 4.4 Å (Figures 2B&C; Suppl. Figure 3B; Table 1). The features of this reconstruction remain 9-fold symmetric. Therefore, we refined class 1, the best resolved class containing 84,460 particles, with C9 symmetry to further improve the resolution. (Refinement including both class 1 and 2 resulted in a slightly poorer resolution.) 3D refinement of class 1 with imposed symmetry results in a reconstruction of the TerS ring to an overall resolution of 3.8 Å (Figures 2D&E; Suppl. Figure 3C; Table 1). Subsequent classification steps with and without alignment did not provide any improvement to the overall resolution.

**Table 1.**
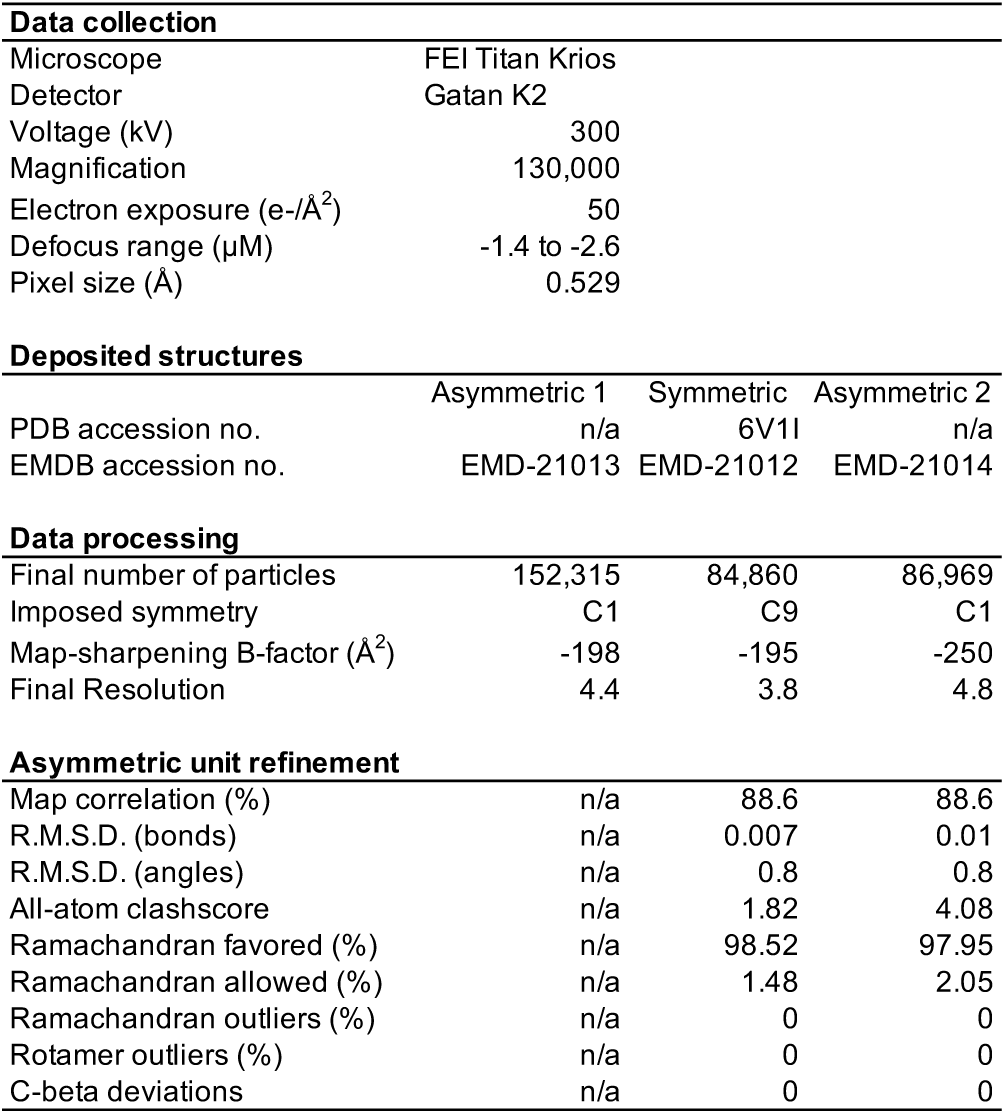
Cryo-EM reconstruction and model refinement statistics.

**Figure 2.**
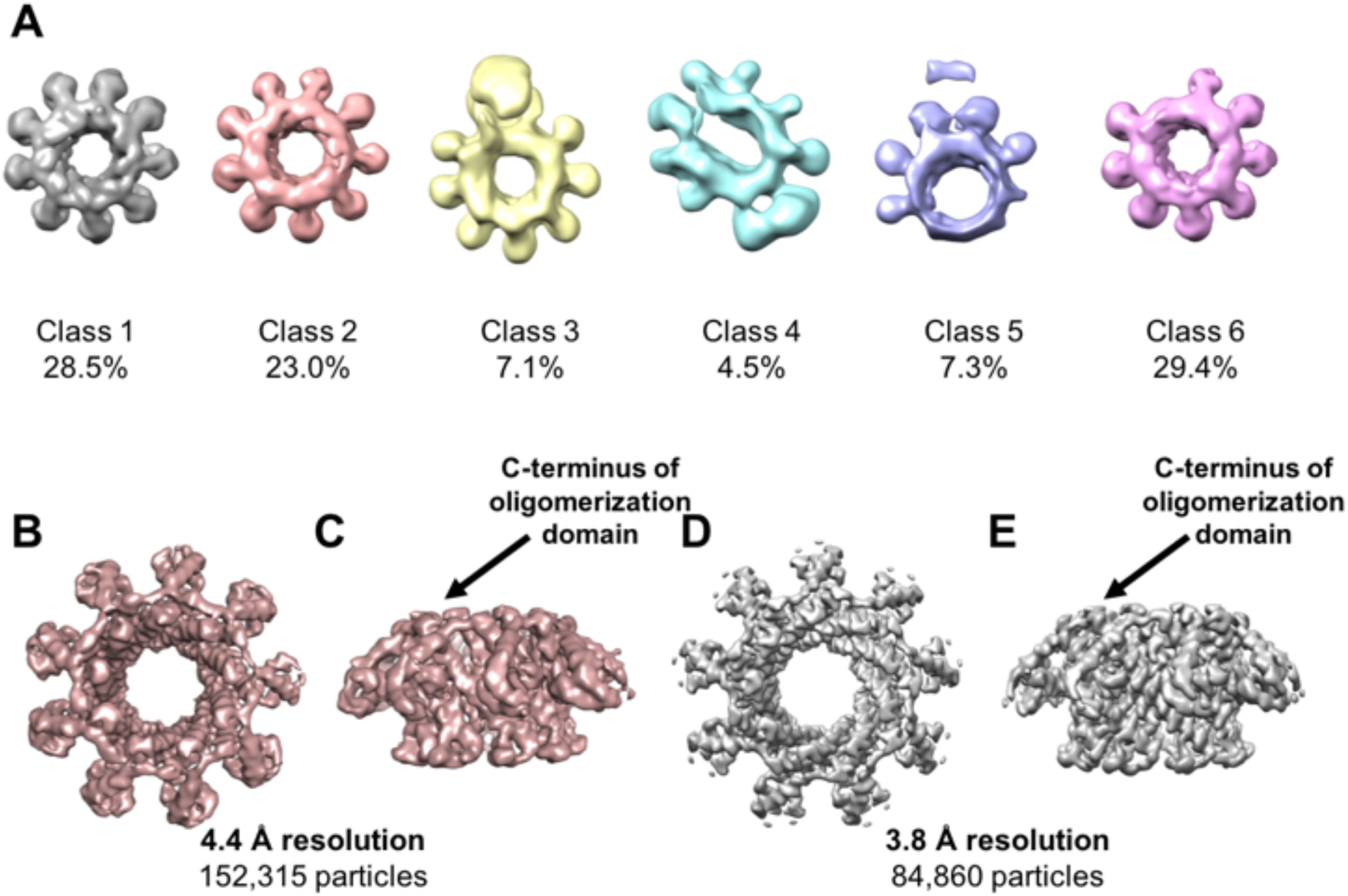
3D Cryo-EM reconstruction of TerS^P74-26^. **(A)** Asymmetric 3D classification shows 9-fold symmetry in the TerS^P74-26^ ring **(B)** 4.4 Å resolution asymmetric 3D reconstruction of the TerS^P74-26^ ring (top). **(C)** Side view of asymmetric TerS reconstruction. **(D)** 3.8 Å resolution C9 symmetric 3D reconstruction of the TerS^P74-26^ ring (top). **(E)** Side view of symmetric TerS reconstruction.

Using the symmetric reconstruction, we built an atomic model of TerS^P74-26^ (Figures 3A-C; Table 1). The model was constructed using the crystal structure of TerS from phage g20c as a starting model (PDB: 4XVN; 98.2% identity to TerS^P74-26^ for the full-length protein). Each TerS^P74-26^ monomer has an N-terminal helix-turn-helix (HTH) motif, followed by an oligomerization domain consisting of two antiparallel helices. These helices pack against the oligomerization domain helices of the neighboring subunit, forming a helical barrel. From the oligomerization domain barrel, the HTH domains extend outward like the spokes of a wheel. The helical barrel arrangement of the oligomerization domains is highly reminiscent of the central oligomerization domains of the TerS proteins from phages SF6 and 44RR, with α-helix 5 of the oligomerization domain positioned in the crevice between α-helices 4 and 5 of the counter-clockwise adjacent subunit when viewed from the C-terminal region (Buttner et al., 2012; Sun et al., 2012) (Figure 3D). The central oligomerization domains appear to be well-ordered, as local resolution of the 3D reconstruction shows the center of the pore has the highest resolution at 3.6 Å (Suppl. Figures 4A&B). The poorest resolution, as low as 4.5 Å, is found around the perimeter of the ring in the tips of HTH domains (Suppl. Figures 4A&B).

**Figure 3.**
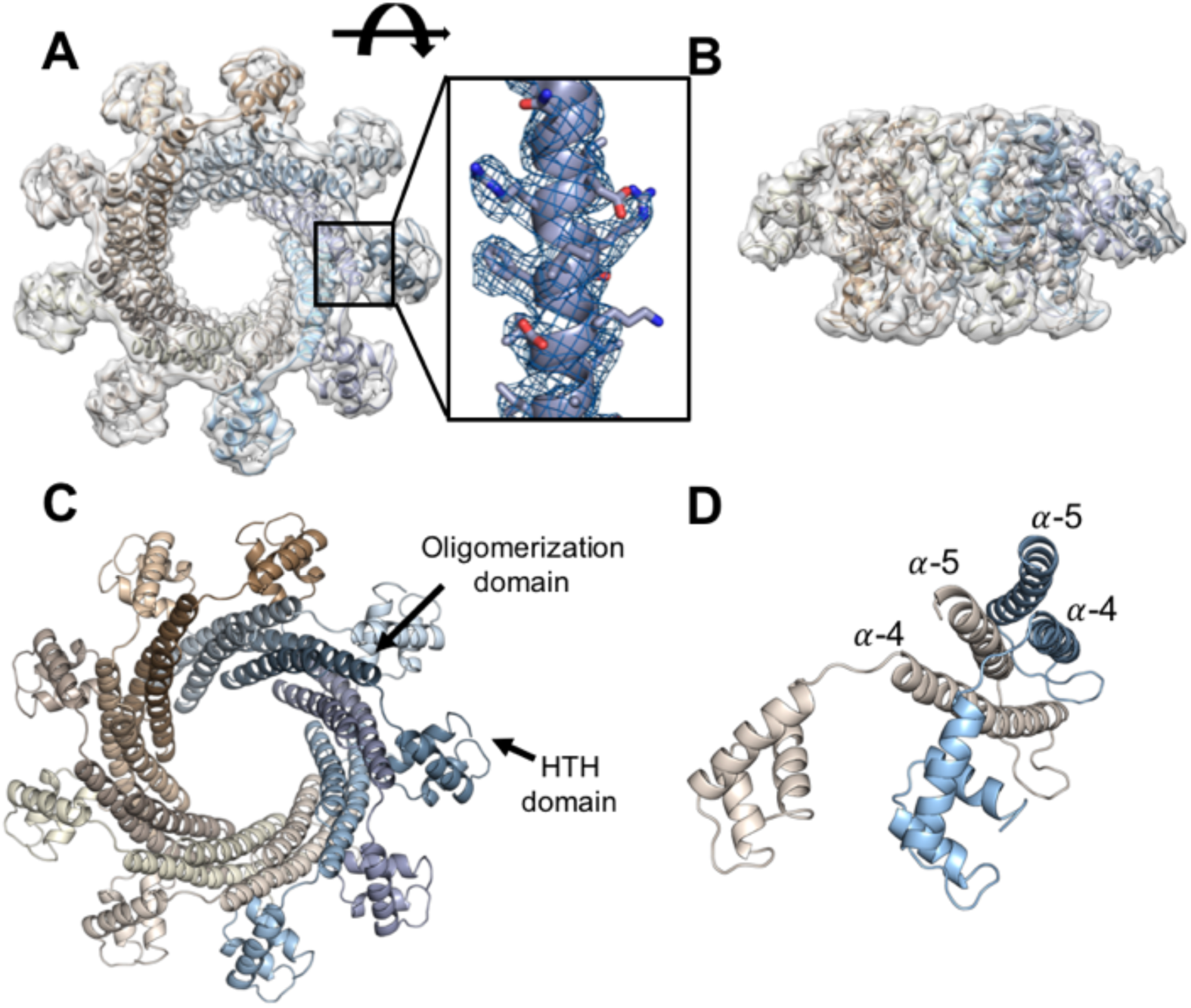
Model of TerS^P74-26^. **(A)** Built atomic model in 3.8 Å resolution TerS^P74-26^ symmetric reconstruction (top). *Inset*: Model built into density of oligomerization domain. **(B)** Side view of atomic model in TerS^P74-26^ reconstruction. **(C)** Top view of atomic model, with HTH and oligomerization domains indicated. **(D**) In each subunit, α-helix 5 packs into the crevice formed by α-helices 4 and 5 in the counter-clockwise subunit. For simplicity, only two subunits (tan and light blue) are shown.

The HTH domain of one subunit interacts with both of the subunits to the right through a series of hydrophobic interactions (Figure 4A). Furthermore, the linker connecting the HTH to the ring (residues 51 to 56) is firmly packed against the adjacent subunit’s oligomerization domain (Figure 4A). Altogether, the HTH domains and linkers bury ∼1570 Å^2^ of area and complete the hydrophobic core of the oligomerization domain. These interactions lock the HTH domains in place, as well as strengthen the nonameric ring by an estimated ∼9 kcal/mol using the PISA server estimation tool (Krissinel and Henrick, 2007). In comparison to mesophilic TerS structures, no other TerS assembly has a similar interaction between the HTH domain and the neighboring oligomerization domain (Fig. 4). We propose that this unique arrangement in TerS^P74-26^ contributes to the rigidification of the HTH domains with implications for DNA binding mechanism (see below).

**Figure 4.**
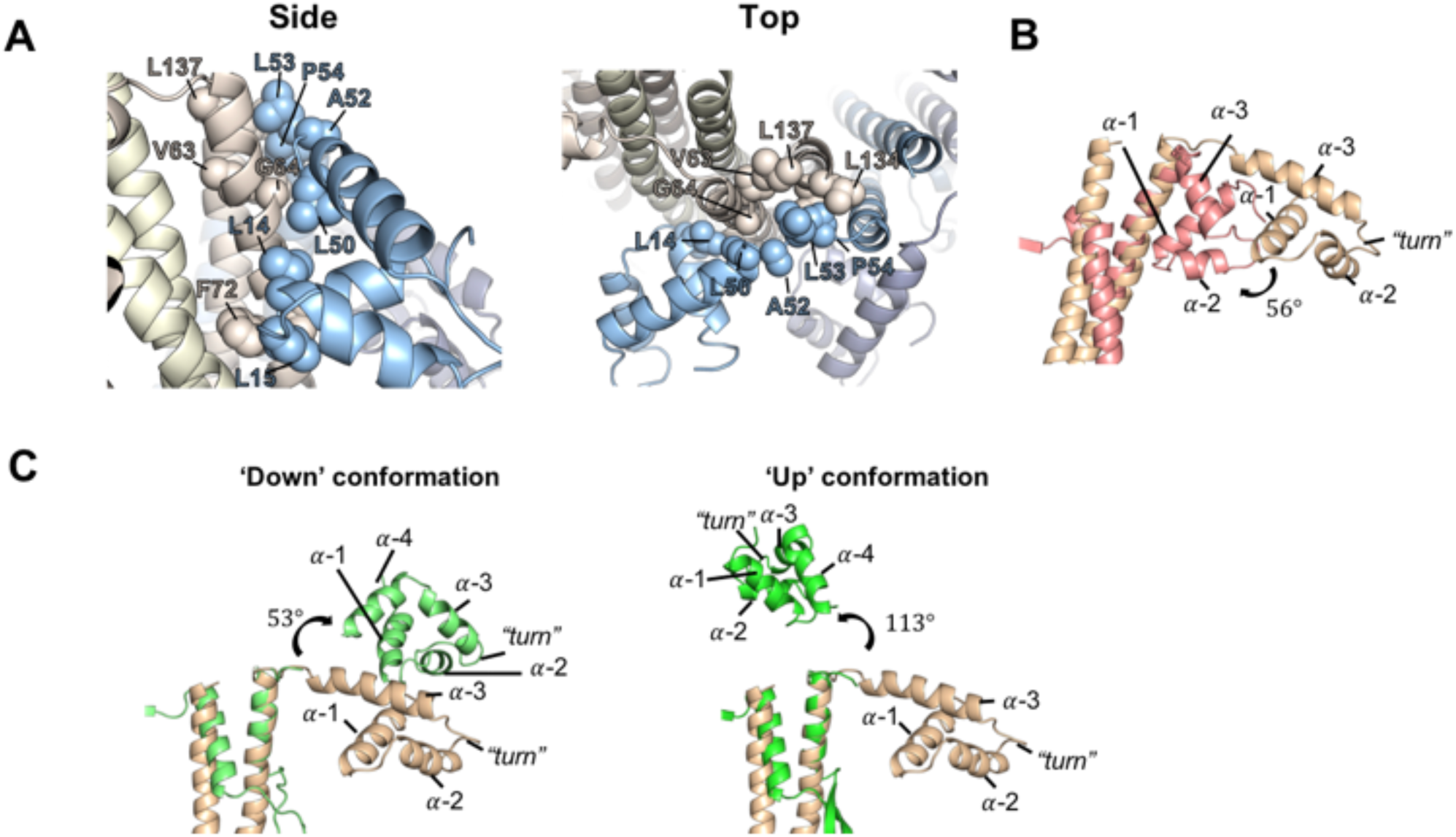
The TerS^P74-26^ linker plays an important role in subunit oligomerization and positions the HTH domain differently than mesophilic TerS proteins. **(A)** Hydrophobic residues (labeled) line the linker (residues 51 to 56) and HTH-oligomerization interfaces between subunits, forming a strong hydrophobic core. **(B)** Alignment of the symmetric TerS^P74-26^ model (tan) with TerS^Sf6^ (pink; PDB 3HEF) shows the TerS^Sf6^ HTH domain is rotated 56° in relation to the TerS^P74-26^ HTH domain **(C)** *Left*: Alignment of the symmetric TerS^P74-26^ model (tan) with oligomerization domain of TerS^SF6^ chain A (light green; PDB 3ZQQ) shows the TerS^SF6^ ‘down’-positioned HTH rotates 53° relative to the TerS^P74-26^ HTH domain. *Right*: Alignment of the symmetric TerS^P74-26^ model (tan) with TerS^SF6^ chain C (green; PDB 3ZQQ) shows the TerS^SF6^ ‘up’-positioned HTH rotates 113° relative to the TerS^P74-26^ HTH domain.

Contrary to our expectations, the last 35 C-terminal residues of the protein are missing in the reconstruction. In mesophilic TerS proteins, this region forms a β-barrel with neighboring subunits and is responsible for TerL binding (Buttner et al., 2012; Gao and Rao, 2011; Roy et al., 2012; Zhao et al., 2010). Both asymmetric and symmetric TerS reconstructions lack density for this region (Figures 2C&E). In 2D classification, side views of the protein show blurry density in the region where the C-terminal region is expected, indicating the region is present, but not resolvable (Suppl. Figure 3A). Interestingly, secondary structure prediction designates this region of TerS^P74-26^ as α-helical (Suppl. Figure 5), which is unexpected because all other TerS structures exhibit C-terminal β-barrels (Buttner et al., 2012; Roy et al., 2012; Zhao et al., 2010).

### Comparison of TerS^P74-26^ with mesophilic TerS proteins

The oligomerization domain of TerS^P74-26^ is similar to that of phage 44RR, a close relative of T4 phage (Sun et al., 2012). In both species, the oligomerization domain consists of two straight, antiparallel helices that assemble into a helical barrel structure (Suppl. Figure 6). The overall C_α_ RMSD of the helices of the oligomerization domains of 44RR and P74-26 is 2.6 Å, suggesting the two domains have considerable structural similarity. However, the barrel of TerS^P74-26^ is a strict 9-mer (Fig 1B), while that of TerS^44RR^ is less well-defined, ranging from an 11-mer to a 12-mer (van Duijn, 2010; Sun et al., 2012). This suggests ring stoichiometry is controlled by slight differences in intersubunit interactions rather than overall secondary structure. In comparison, TerS of *Shigella* phage Sf6 uses a similar fold of antiparallel helices, although the helices are quite bent (Suppl. Figure 6) (Zhao et al., 2010). Furthermore, the interactions between neighboring oligomerization domains of TerS^Sf6^ are different than in other TerS proteins, as was pointed out previously (Sun et al., 2012). The oligomerization domain of TerS of *Bacillus* phage SF6 is also quite distinct, with a beta-hairpin inserted at the turn between the two antiparallel helices; these twisted beta-hairpins extend the barrel structure formed by the helical region of the oligomerization domain (Buttner et al., 2012). Despite the substantial differences in primary amino acid sequence, secondary structure (Suppl. Figure 7), and mechanism for assembly, the overall structure is remarkably similar across phage, with barrel architecture retaining an overall outer dimension of 52 to 77 Å between C_α_ atoms across the barrel. Therefore, we hypothesize that the core oligomerization fold is not conserved, but rather a barrel shape

The HTH domain of TerS^P74-26^ is also arranged distinctly from other phage. In TerS^SF6^ and TerS^Sf6^, the HTH domains are flexible in regards to the central oligomerization domain (Buttner et al., 2012; Zhao et al., 2010, 2012). It is speculated that this flexibility permits the HTH domains to stagger during DNA wrapping, allowing DNA to adopt a less strained conformation. We performed several analyses to investigate if the same conformational changes occur between the HTH domains of TerS^P74-26^. First, we examined class 6 (86,969 particles), which is the most asymmetric class, with only eight HTH domains visible (Figure 2A). As other TerS structures show flexibility in the HTH domains (Buttner et al., 2012; Zhao et al., 2010, 2012), it is possible the missing domain in this class is due to the inherent flexibility of this region. 3D refinement with no symmetry applied produces a reconstruction with an overall resolution of 4.8 Å (Suppl. Figure 8A-C; Table 1). The reconstruction was used to create an atomic model of the class 6 structure by rigid body fitting each domain of the symmetrical model into the density (Suppl. Figure 8D; Table 1). Comparing each chain of the class 6 asymmetric model to all other chains within the model, no differences in HTH motif orientation relative to the oligomerization domains were observed (Suppl. Figure 8E). To determine if the missing HTH domain is the result of proteolytic removal rather than protein flexibility, we ran concentrated purified protein on an SDS-PAGE gel. The gel shows minor proteolysis of TerS, with a band at the approximate size of a subunit missing a HTH domain (Suppl. Figure 8F). Using gel densitometry, we estimate that approximately 4.5% of the protein is proteolysed, which is comparable to the ∼3% estimated by cryo-EM. This result suggests that the missing HTH domain in class 6 is due to proteolysis, rather than conformational heterogeneity within the TerS ring. Our attempts to visualize any conformational heterogeneity using multi-body refinement or localized reconstruction methods were complicated by the small size of the HTH domain (∼6 kDa; data not shown). Nonetheless, our data indicate very little conformational heterogeneity in the HTH domains of TerS^P74-26^.

The arrangement of the HTH domains around the perimeter of the ring is critical for examining the wrapping model that has been proposed for most TerS proteins (Buttner et al., 2012; Gao et al., 2016; Zhao et al., 2010, 2012). HTH domains usually contain three helices and interact with the DNA major groove using α-helix 3 (Aravind et al., 2005). In comparison to the crystal structure of *Shigella* phage Sf6 TerS, the P74-26 HTH domains extend outward and rotate 56° counter-clockwise with respect to the central oligomerization domains (Figure 4B). This rotation positions α-helix 3 of TerS^P74-26^ nearly perpendicular to the central oligomerization domains, whereas in Sf6 this helix is at a 70° angle relative to the oligomerization domains. In the crystal structure of *Bacillus* SF6 TerS, the three HTH domains in the asymmetric unit are tethered to the ring by highly flexible linkers, with one HTH domain invisible and the other two positioned in dramatically different orientations (Buttner et al., 2012). Neither of the two visible conformations of TerS^SF6^ are similar to that observed in TerS^P74-26^. While one HTH domain of TerS^SF6^ is oriented downward similarly to TerS^P74-26^, it exhibits a 53° clockwise rotation with respect to the oligomerization domain (Figure 4C). The second HTH orientation in the SF6 crystal structure is even more dissimilar, and is positioned in an ‘up’ conformation with a 113° clockwise rotation (Figure 4C). Therefore, in comparison to Sf6 and SF6 TerS proteins, the helix-turn-helix domains of the P74-26 TerS model are oriented differently in relation to the oligomerization domains, suggesting there are mechanistic distinctions in how the three TerS proteins bind DNA.

The ‘turn’ of the HTH domain in TerS^P74-26^ contains basic and polar residues. These residues, specifically Lys31, Arg32, Lys33, and Thr35, may potentially bind the DNA phosphate backbone. In phage SF6, it was shown that residues in his ‘turn’ region confer a non-specific effect on DNA binding (Greive et al., 2016). Helix 3 of TerS^P74-26^ is also lined with polar and charged residues (Suppl. Fig 9). This is similar to that found in other HTH domains (Beamer and Pabo, 1991; Brennan et al., 1990; Schultz et al., 1991). From this, we predict that the ‘turn’ region of the P74-26 HTH domain primarily binds DNA phosphates through non-specific interactions, while polar residues of helix 3 interact with DNA bases and sugars.

## DISCUSSION

### The unresolved C-terminal region

A C-terminal β-barrel region is thought to be a necessary component in other phage TerS proteins, as the β-barrel stabilizes the oligomerization state of the complex and its removal results in polydisperse oligomers (Buttner et al., 2012; Sun et al., 2012). The formation of the barrel requires strict interactions between β-strands of neighboring subunits, which enforces proper stoichiometry of the ring. However, in our extensive analysis of the cryo-EM data, we find no evidence of β-barrel formation, yet our TerS assemblies remain completely monodisperse according to SEC-MALS (Figure 1B). Moreover, the crystal structure of the nearly identical TerS protein from the Antson Lab (PDB code 4XVN) also lacks any density for this region. Therefore, we propose that a β-barrel is not critical for retaining correct stoichiometry in TerS^P74-26^.

Additionally, it is known that the TerS C-terminal region makes critical contacts with the large terminase for packaging (Gao and Rao, 2011; Roy et al., 2012). This raises the question of how the small terminase of this thermophilic phage binds TerL, and what the nature of this interaction is. It is possible that TerS^P74-26^ requires a partner, such as DNA, TerL, or a different protein to order the C-terminal region. Because the C-terminal region is predicted to be alpha-helical, this interaction mechanism could be distinct from that of TerS proteins from other phage with β-barrel domains. The lack of a rigid connection between the β-barrel and the oligomerization domain core could have a functional role, as perhaps this flexibility allows the motor to function more efficiently. Future studies will investigate this issue.

### The role of the fixed HTH domains in binding DNA

The HTH domains of TerS^P74-26^ are rigidly bound to the central hub of oligomerization domains. This is in contrast to structures of TerS rings from other phage in which the HTH domains are flexibly tethered to the hub. The interaction between the HTH domains and the oligomerization hub is mediated by residues in the cleft between helices 1 and 3 of the HTH domain. Other HTH domains often have idiosyncratic interactions or structural features that are positioned within this cleft, suggesting that the cleft is a hotspot for evolution of new interactions (Aravind 2005 FEMS Microbiology). We hypothesize that this interface was evolved by progenitors of TerS^P74-26^ to increase the stability of the TerS ring. By tightly locking to the oligomerization domains of neighboring subunits, the HTH domains of TerS^P74-26^ can also play a role in stabilizing the overall ring assembly. The interface formed between HTH domains and the neighboring oligomerization domains is substantial, and consists primarily of hydrophobic interactions (Figure 4A). Because the entropically-driven hydrophobic effect becomes stronger at increasing temperature (Huang and Chandler, 2000), we anticipate the HTH domains remain locked in place even at the elevated temperature environment of phage P74-26.

This unique interaction between the HTH and oligomerization domain serves to enforce the stability and stoichiometry of the TerS^P74-26^ ring. The linker between the HTH and oligomerization domain is nearly fully extended, yet locked in place through hydrophobic interactions forming part of the hydrophobic core (Figure 4A). This constrains ring stoichiometry, as each HTH domain contacts two other subunits within the assembly through this linker, and other oligomeric states would likely not support the geometry of these interactions. With strict HTH-oligomerization domain interactions enforcing stability and stoichiometry of the ring, the constraints of an ordered β-barrel domain are released. Thus, we hypothesize that these interactions allowed the C-terminal β-barrel domain of TerS^P74-26^ to no longer adopt a rigid conformation relative to the oligomerization domain.

Furthermore, we propose the conformation of the HTH domains observed for apo-TerS^P74-26^ represents the overall location and orientation of TerS HTH motifs after DNA binding. Although we currently lack a DNA-bound structure of TerS^P74-26^, the tight interaction between HTH and oligomerization domains makes it doubtful that the ring undergoes a substantial rearrangement upon binding DNA. If the HTH domain releases from the oligomerization domain, this would expose the hydrophobic core and linker to solvent. The energetic penalty for hydrophobic exposure would be even more acute at the elevated temperature of P74-26’s native environment. Therefore, it is likely that the HTH domains remain locked into position, even after DNA binding.

The fixed orientation of the HTH domains places major constraints on how TerS^P74-26^ wraps DNA around the ring. HTH domains most often bind DNA by inserting the recognition helix (in ringed TerS proteins helix 3) into the DNA major groove to achieve specificity, with residues in the ‘turn’ used for binding the phosphate backbone (Beamer and Pabo, 1991; Brennan et al., 1990; Schultz et al., 1991). The homologous protein TerS^SF6^ appears to adopt this typical HTH-DNA binding mode, as the ‘turn’ and N-terminal region of α-helix 3 contributes to non-specific DNA binding (Greive et al., 2016). In TerS^P74-26^, the localization of basic residues in this region (Lys31, Arg32, Lys33) creates a positively-charged surface (Figure 5B) that could potentially interact with negatively-charged DNA phosphates. Helix 3 of TerS^P74-26^ lies on the top of the HTH domain, with the exposed surface containing several polar groups that could be used for hydrogen bonding to DNA bases and sugars (Suppl. Figure 9). Therefore, we predict that the DNA is positioned along the ‘top’ of the HTH domains of TerS^P74-26^. The spacing between helix 3 of adjacent subunits is ∼30 Å, which is approximately what is expected for the major groove spacing within DNA wrapping around the TerS^P74-26^ ring (∼80-100 Å diameter between recognition helices). As a point of comparison, the major groove spacing in nucleosomal DNA is slightly tighter (∼28 Å), for wrapping around a particle that is smaller (∼65 Å) (Luger et al., 1997).

**Figure 5.**
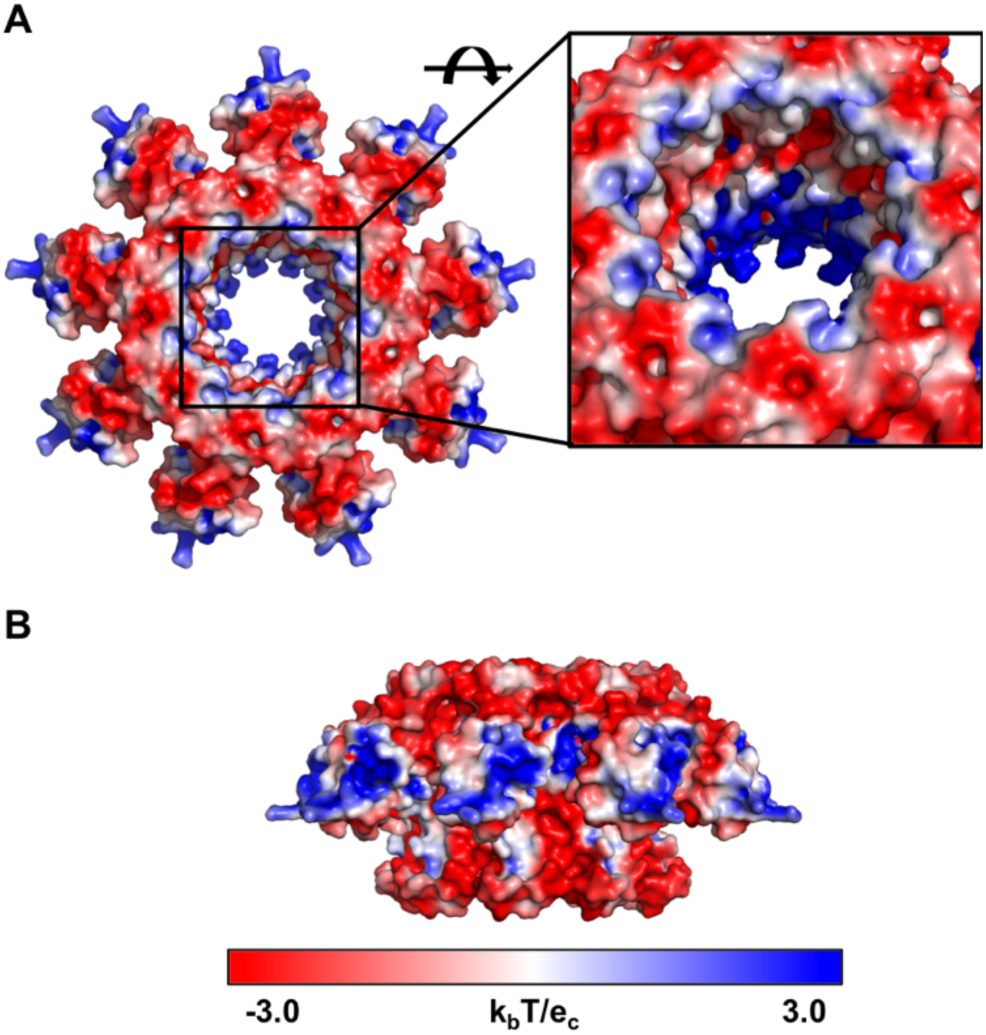
Electrostatics of the TerS^P74-26^ ring. **(A)** Top and **(B)** side views of TerS^P74-26^ electrostatics using the APBS Pymol Plugin (Delano Scientific). Blue coloring indicates a net positive charge, while red coloring indicates a net negative charge. Positive charges are concentrated in the HTH domains and the center of the TerS pore. *Inset:* Negative and positively charged regions alternate within the TerS pore.

**Figure 6.**
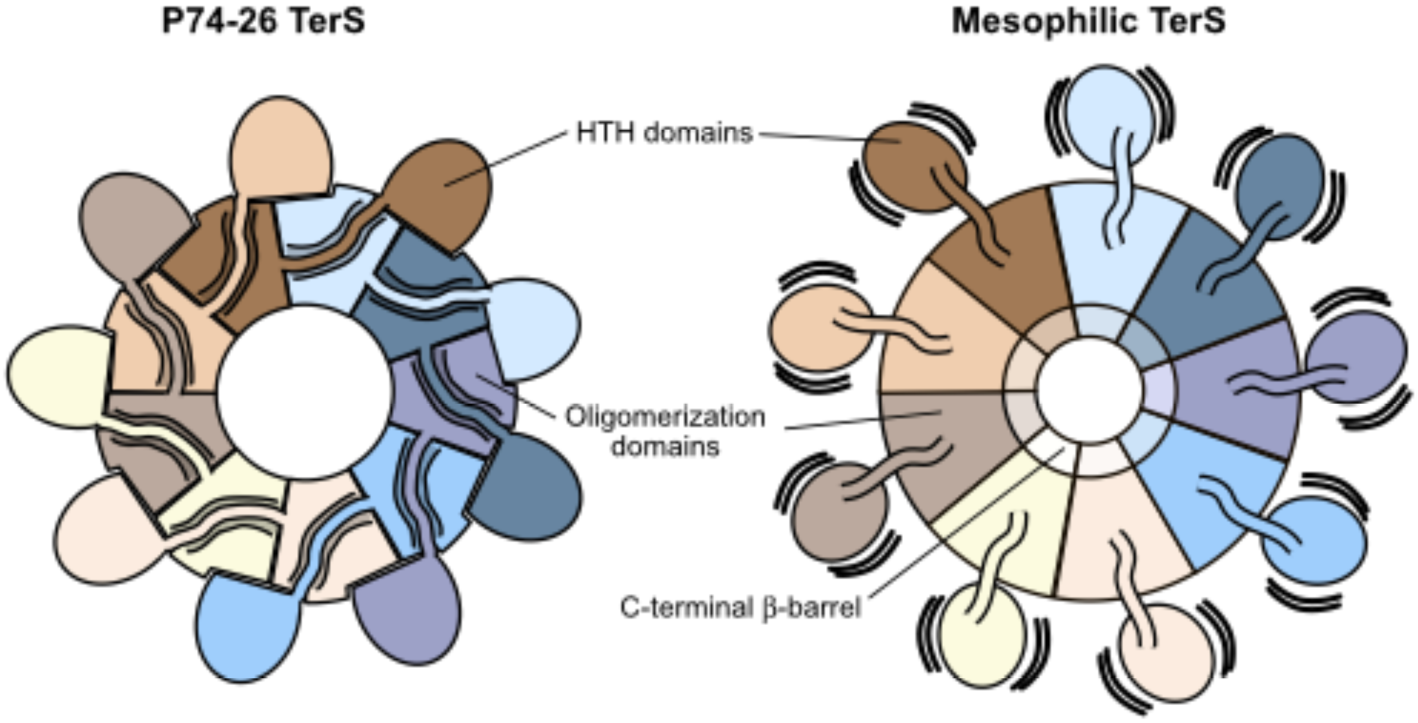
Comparison of TerS^P74-26^ with mesophilic TerS complexes. *Left*: Intersubunit interactions between the HTH domain, domain linker, and neighboring clockwise oligomerization domains lock HTH domains into place in TerS^P74-26^ rings, stabilizing the conformation of the HTH domains. *Right*: In mesophilic TerS assemblies, the HTH domains and domain linkers do not form tight interactions with neighboring oligomerization domains, allowing the HTH domains to adopt flexible conformations in relation to the core assembly.

We hypothesize a different DNA binding mode for TerS^P74-26^ compared to its mesophilic cousins. DNA wrapping would favor superhelix formation, as this allows the two ends of DNA to freely pass each other without steric hindrance (Suppl. Figure 10) (An example of a superhelix would be the nucleosome, in which the DNA spirals around the histone core.) The flexibility in the HTH domains observed for TerS^SF6^ and TerS^Sf6^ could possibly accommodate superhelix formation. However, the rigid orientations of the TerS^P74-26^ HTH domains would prevent a superhelical conformation. Therefore, we propose that at least one of the HTH domains is disengaged from DNA to allow DNA to pass by the other end unimpeded. Future studies will examine how DNA binding and sequence recognition are achieved.

Alternatively, DNA could thread through the central pore instead of wrapping around the HTH domains. The narrowest diameter of the TerS^P74-26^ pore is 29 Å, which is large enough to accommodate double-stranded DNA (∼20 Å diameter). Although some TerS proteins have central pores too small to accept double-stranded DNA (Buttner et al., 2012; Zhao et al., 2010), TerS^P22^ is hypothesized to bind DNA using a threading mechanism, as it lacks a HTH domain (Roy et al., 2012). Interestingly, it’s predicted that TerS^P22^ has an α-helical C-terminal region following the β-barrel (Roy et al., 2012), similar to the secondary structure prediction of TerS^P74-26^ (Suppl. Figure 5; Suppl. Table 1). The inner pore of TerS^P74-26^ has mixed electrostatic surface, with interspersed layers of basic and acidic residues. (Figure 5A). The pore surface may potentially form tracts of attractive and repulsive DNA binding regions. If DNA threads through the central pore, the DNA may tilt relative to the central pore axis of TerS^P74-26^ to avoid interactions with acidic residues. There is precedent for an off-axis mode of DNA binding within a ring, as DNA binds inside DNA polymerase sliding clamps in a tilted fashion (Georgescu et al., 2008; McNally et al., 2010). Future studies will test this threading model. (It is worth mentioning that the threading and wrapping models are not mutually exclusive.)

Together, our work presents a novel thermophilic system for studying small terminase proteins and their role in viral maturation. To our knowledge, this is the first cryo-EM structure of a small terminase protein at a resolution permitting atomic modeling, yet the C-terminal region is not well-ordered. Future studies of TerS^P74-26^ will elucidate the conformation of the C-terminal region and its role in TerL binding and enzymatic regulation, as well as the DNA binding mechanism.

## MATERIALS AND METHODS

### Cloning

The TerS^P74-26^ gene was synthesized with codon optimization for expression in *E. coli* by Genscript Corporation. The gene was cloned into the BamHI and NdeI sites of a modified pET28a vector with an N-terminal His6-T7-gp10 expression tag and a Prescission protease cut site. Enzymes were purchased from New England BioLabs. Oligonucleotides were purchased from IDT.

### Protein expression and purification

Protein was expressed in BL21-DE3 cells containing the pET28a-TerS plasmid. Bacterial cultures were grown at 37°C in Terrific Broth supplemented with 30 µg/ml kanamycin until an OD600 of 0.7 was reached. Cells were moved to 4°C for 20 minutes, after which expression was induced by addition of IPTG (isopropyl - b - D - thiogalactopyranoside) to 1 mM. Cells were then returned to an 18°C incubator to shake overnight. Cells were pelleted and resuspended in ‘Buffer A’ (500 mM NaCl, 20 mM Tris pH 7.5, 20 mM imidazole and Roche cOmplete™ EDTA-free Protease Inhibitor Cocktail dissolved to a final concentration of 1x). Resuspended cells were flash frozen in liquid nitrogen for long-term storage at -80°C. Thawed cells were lysed using a cell disrupter, and lysis was pelleted by centrifugation. Cleared lysate was filtered using a 0.45 µM filter. All subsequent steps occurred at room temperature unless noted. Lysate was loaded and recirculated over Ni-affinity beads (Thermo-Scientific) for 2.5 hours, which had been pre-equilibrated with Buffer A. Beads were subsequently washed with 5 column volumes of Buffer A without protease inhibitors. The protein-bound beads were transferred to a 50 mL conical containing 1.25 mg of purified prescission protease, which was incubated overnight on a nutator. The following day, the resin was transferred to a gravity flow column, and the flow-through was collected, alongside a 1 column volume wash of the resin with Buffer A. The flow-through was then concentrated and injected onto a HiPrep 26/60 Sephacryl S200-HR gel filtration column that had been pre-equilibrated with gel filtration buffer (250 mM NaCl, 20 mM Tris pH 7.5) at 4°C. Fractions corresponding to the TerS peak were pooled, concentrated to 17 mg/mL, and flash frozen in liquid nitrogen for storage at -80°C. TerL^P74-26^ was expressed and purified as previously described (Hilbert et al., 2015).

### Size exclusion chromatography Multi-angle light scattering (SEC-MALS)

SEC-MALS was performed at room temperature using a 1260 Infinity HPLC system (Agilent), a Dawn Helios-II multi-angle light scattering detector (Wyatt Technology), and an Optilab T-rEX differential refractive index detector (Wyatt Technology). Detectors were aligned, corrected for band broadening, and photodiodes were normalized using a BSA standard. Samples were diluted to 1 mg/mL with Gel Filtration buffer and filtered through a 0.22 µM filter. 50 µL of sample was injected onto a WTC-030S5 size exclusion column with a guard (Wyatt Technology) that had been pre-equilibrated overnight with Gel Filtration buffer. Data analysis was performed with Astra 6 software (Wyatt Technology).

### DNA binding and enzymatic assays

TerS DNA binding was performed using the P74-26 gp83 DNA sequence that was PCR amplified from the P74-26 phage genome. P74 - 26 forward primer: ATGAGCGTGAGTTTTAGGGACAGGG; P74-26 reverse primer: CTAGGTC TTAGGCGTTTCATCCGCC. Oligonucleotides were purchased from IDT. To assess DNA binding, TerS was dialyzed into a buffer containing 25 mM potassium glutamate and 10 mM Tris pH 7.5. TerS was then incubated for 30 minutes with 50 ng of the P74-26 gp83 gene in an 8 µL volume sample. After incubation, 2 µL of 5x Orange G loading dye was added to the samples, yielding the final protein concentration indicated on the gel. Samples were run on a 1% (wt/vol) TAE-agarose gel with a 1:10,000 dilution of GelRed dye (Phenix Research) for 90 minutes at 80 volts. ATPase and nuclease experiments were performed as previously described (Hilbert et al., 2015, 2017).

### Electron Microscopy

#### Negative Stain EM

3.5 µL of 900 nM TerS (monomer) was applied to a glow-discharged carbon-coated 400 mesh copper EM grid and incubated for 30 seconds. Sample was blotted off, and the grid was washed with water and blotted two times. Grid was stained with 1% uranyl acetate and imaged using a 120kV Philips CM-120 electron microscope with a Gatan Orius SC1000 detector. Relion 2.0 was used for 2D classification (Kimanius et al., 2016).

#### Cryo-EM sample preparation

For dataset one, 400 mesh 2/2 Holey Carbon C-Flat grids (Protochips) were incubated with ethyl acetate until dry. Grids were glow discharged for 60 seconds at 20 mA (negative polarity) with a Pelco easiGlow glow discharge system (Pelco). Samples were prepared to yield a final concentration of 19.5 µM TerS (nonamer), 150 mM NaCl, 20 mM Tris (pH 7.5), and 0.015% amphipol A8-35. For dataset two, the same sample was applied to a 200 mesh 2/2 UltrAuFoil Holey Gold grid (Quantifoil) that was glow-discharged for 60 seconds at 20 mA. For both datasets, 3 µL of sample was applied to the grid at 10°C and 95% humidity in a Vitrobot Mark IV (FEI). Samples were blotted for 4 seconds with a blot force of 5 after a 10 second wait time. Samples were then vitrified by plunging into liquid ethane and were stored in liquid nitrogen until data collection.

#### Cryo-EM data collection

Micrographs were collected on a Titan Krios electron microscope (FEI) at 300 kV fitted with a K2 Summit direct electron detector (Gatan). Images were collected at 130,000x in superresolution mode with a pixel size of 0.529 Å/pixel and a total dose of 50 e-/Å2 per micrograph. Micrographs were collected with a target defocus range of -1.4 to -2.6 for both datasets one and two. Dataset one was collected with one shot focused on the center of the hole. For dataset two, the first 549 images were collected with four shots per hole at 0° tilt, and the remaining 1,077 images were collected at a 30° tilt with two shots per hole. After combining datasets 1 and 2, a total of 2,822 micrographs were collected.

#### Data Processing

Micrograph frames were aligned using the Align Frames module in IMOD with 2x binning, resulting in a final pixel size of 1.059 Å/pixel. Initial CTF estimation was performed using CTFFIND (Rohou and Grigorieff, 2015) within the cisTEM suite. Particles were picked with a characteristic radius of 40 Å using ‘Find Particles’ in the cisTEM software package (Grant et al., 2018). Particles were then extracted with a largest dimension of 120 Å and a box size of 256 pixels. Selected particles were subjected to 7 rounds of 2D classification using cisTEM. Each round of 2D classification consisted of 20 iterative cycles with 50 to 100 classes. After each round, the classes were examined and noisy classes were excluded before subjection to the next round of classification. The final round of 2D classification yielded 295,395 particles, which were exported into Relion format.

Ab-initio 3D reconstruction was performed with cisTEM using a particle subset selected for an even distribution of views from the 2D classification images. Ab-initio 3D reconstruction was performed using 2 starts with 40 cycles per start. CTF correction was re-estimated using gCFT (Zhang, 2016) and the particles were re-extracted in Relion 3.0 (Zivanov et al.). 3D Classification was done in Relion 3.0 using C1 symmetry into 6 classes for 60 iterations with a mask diameter of 140 Å. For the first asymmetric structure, classes 1 and 2 were combined (152,315 particles) for 3D refinement in Relion 3.0 using C1 symmetry. For the symmetric reconstruction, class 1 (84,860 particles) was sub-selected for 3D refinement in Relion 3.0 using C9 symmetry. For the second asymmetric structure, class 6 (86,969 particles) was sub-selected for asymmetric refinement using C1 symmetry. CTF refinement and subsequent post-processing were performed after 3D refinement for all symmetric and asymmetric reconstructions in Relion 3.0. Resolution was calculated using gold-standard FSC curve calculation and a cutoff of 0.143.

#### Model Building

To build the atomic models of the TerS structure, the helix-turn-helix motifs and oligomerization domains of the g20c crystal structure (PDB code 4XVN) were rigid body fit into the cryo-EM density for each subunit separately using the Chimera ‘Fit to map’ command (Pettersen et al., 2004). Each chain in the symmetric and asymmetric models consisted of residues 1 to 137. For the symmetric structure, one chain was manually refined in Coot (Emsley et al., 2010), and 9-fold symmetry was repopulated using PyMol. For the class 6 asymmetric structure, the symmetric model was fit into the density and each helix-turn-helix motif and oligomerization domain were separately fit in Coot using the ‘rigid body refine’ tool. Model refinement was performed in Phenix using the real-space refinement tool with three cycles of refinement per round. Rotamer restraints, Ramachandran restraints, and NCS restraints were used during refinement. Group ADP values were calculated on a per residue basis. Electrostatic maps were generated using the PyMol APBS plugin.

### Secondary Structure Analysis

Secondary structure of TerS monomers was predicted using JPred4 (Drozdetskiy et al., 2015). Structure-based sequence alignments were performed with ‘chain A’ of the PDB structures 3ZQQ, 3HEF, and 3TXQ using PROMALS3D (Pei et al., 2008). Structure alignment figure was created using ESPript 3 (Robert and Gouet, 2014).

EMDB Accession codes are EMD-21012, EMD-21013, EMD-21014. PDB accession code is 6V1I.

## Author Contributions and Notes

JAH and BAK designed research, JAH and BJH performed research, JAH, BJH, C.G., N.P.S., and BAK analyzed data; and JAH and BAK wrote the paper. CG and NPS provided valuable insight into optimization of cryo samples and reconstruction refinement.

The authors declare no conflict of interest.

This article contains supporting information online.

## Acknowledgments

The authors thank Drs. C. Xu, KK Song, and K. Lee for assistance with data collection, and Drs. C. Xu, A. Korostelev, and Mrs. A. Jecrois for advice on data processing. We thank members of the Kelch, Royer and Schiffer labs for helpful discussions. JAH was supported by an NRSA Predoctoral Fellowship from NIGMS (1F31GM121019-01A1). This work was funded by the Pew Charitable Trusts and the National Science Foundation (1817338).

