## Supplemental Material for "A thermophilic phage uses a small terminase protein with a fixed helix-turn-helix geometry"

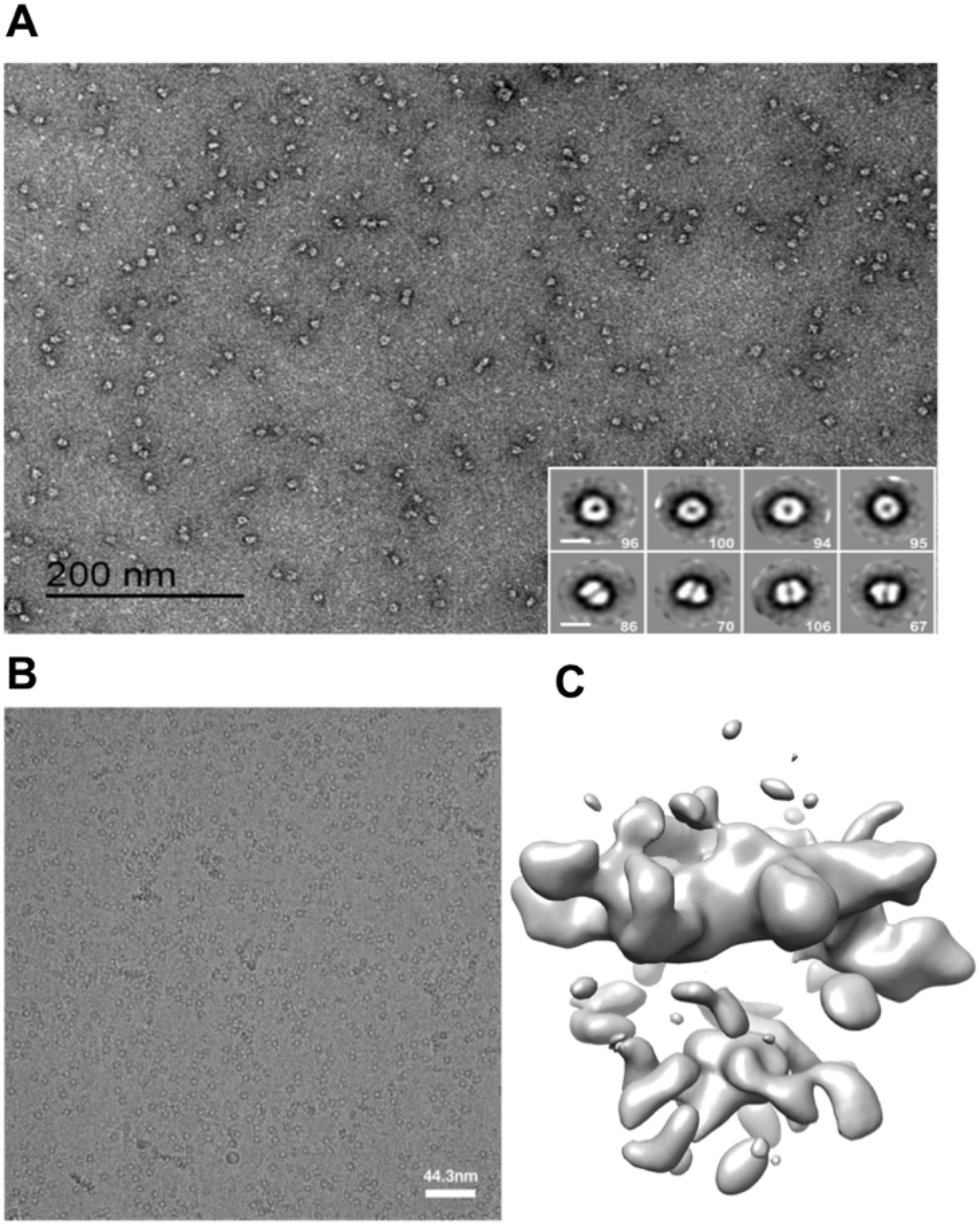


**Supplemental Figure 1. Preliminary electron microscopy analysis of TerS$^{P74-26}$**. **(A)** Negative stain of TerS$^{P74-26}$ shows that particles are homogenous and evenly dispersed. 2D classification (*inset*) shows an even distribution of top and side views. **(B)** Cryo-EM micrograph shows mostly top and bottom views of the TerS$^{P74-26}$ ring. Image taken on FEI Talos Arctica at 200 kV with a Gatan K2 Direct Detector. **(C)** 3D reconstruction (side view) lacks density in center due to preferred orientation.

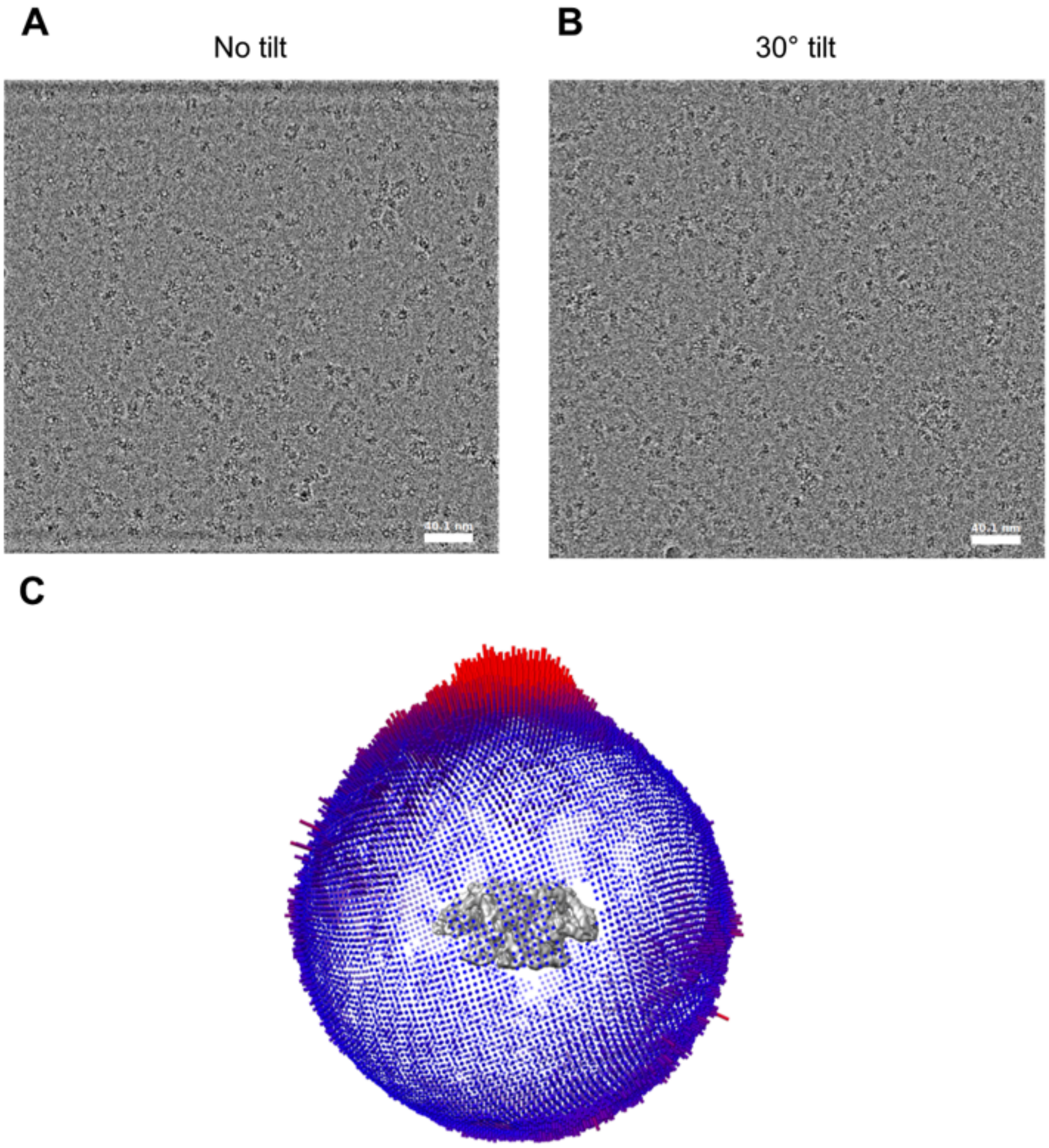


**Supplemental Figure 2. Tilted and un-tilted data collection on gold grids.** Images taken on Thermo Fisher Scientific Titan Krios with a Gatan K2 Direct Detector. **(A)** un-tilted and **(B)** 30° tilted micrographs were used to provide side views of the TerS$^{P74-26}$ complex. **(C)** Angular distribution of particles for asymmetric reconstruction of classes 1 and 2. The TerS$^{P74-26}$ particles still show preferred orientation, but enough side views are captured for 3D reconstruction.

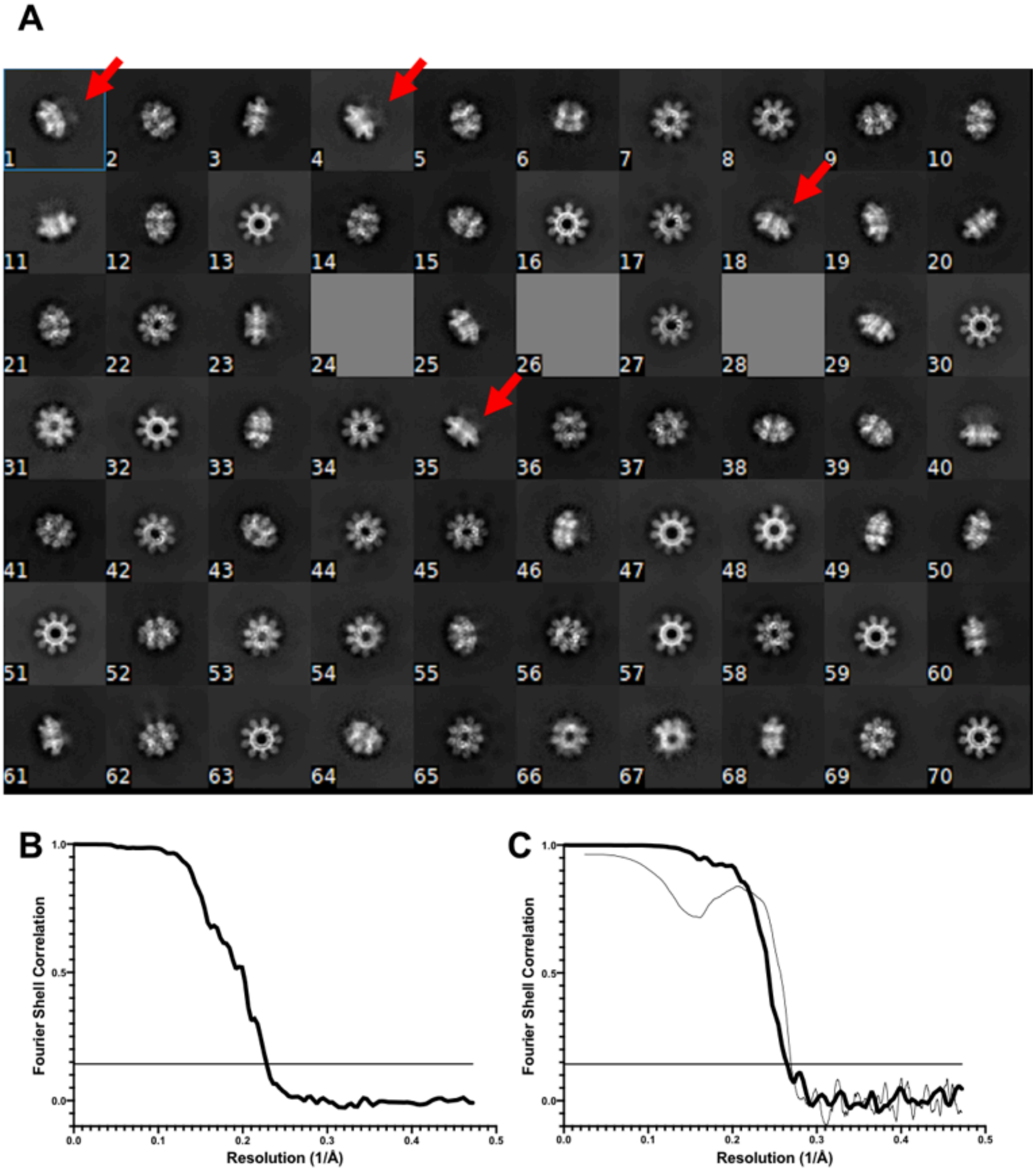


**Supplemental Figure 3. TerS$^{P74-26}$ cryo-EM structure determination. (A)** TerS$^{P74-26}$ 2D classification shows blurred density where C-terminal region should be (red arrows). **(B)** Gold-standard FSC curve of class 1 and 2 asymmetric 3D reconstruction. Flat line represents the FSC 0.143 cutoff, indicating an overall resolution of 4.4 Å. **(C)** Gold-standard FSC curve of symmetric class 1 3D reconstruction (thick black line) and model to map FSC (thin black line). Flat line represents the FSC 0.143 cutoff, indicating an overall resolution of 3.8 Å.

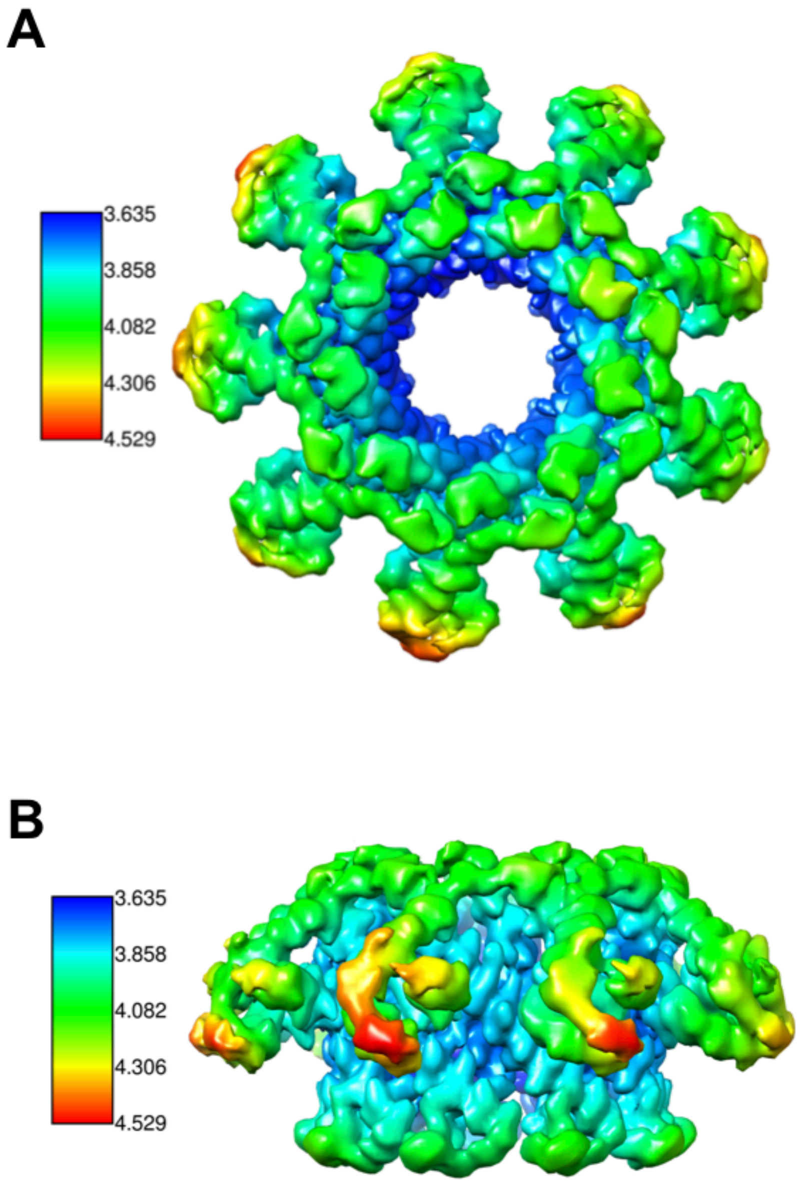


**Supplemental Figure 4. Local resolution of the TerS$^{P74-26}$ ring**. Highest resolution of the TerS$^{P74-26}$ ring is in the center of the pore, while the lowest resolution is in the ‘turn’ region of the helix-turn-helix domain. **(A)** Top view of the ring **(B)** Side view of the ring.

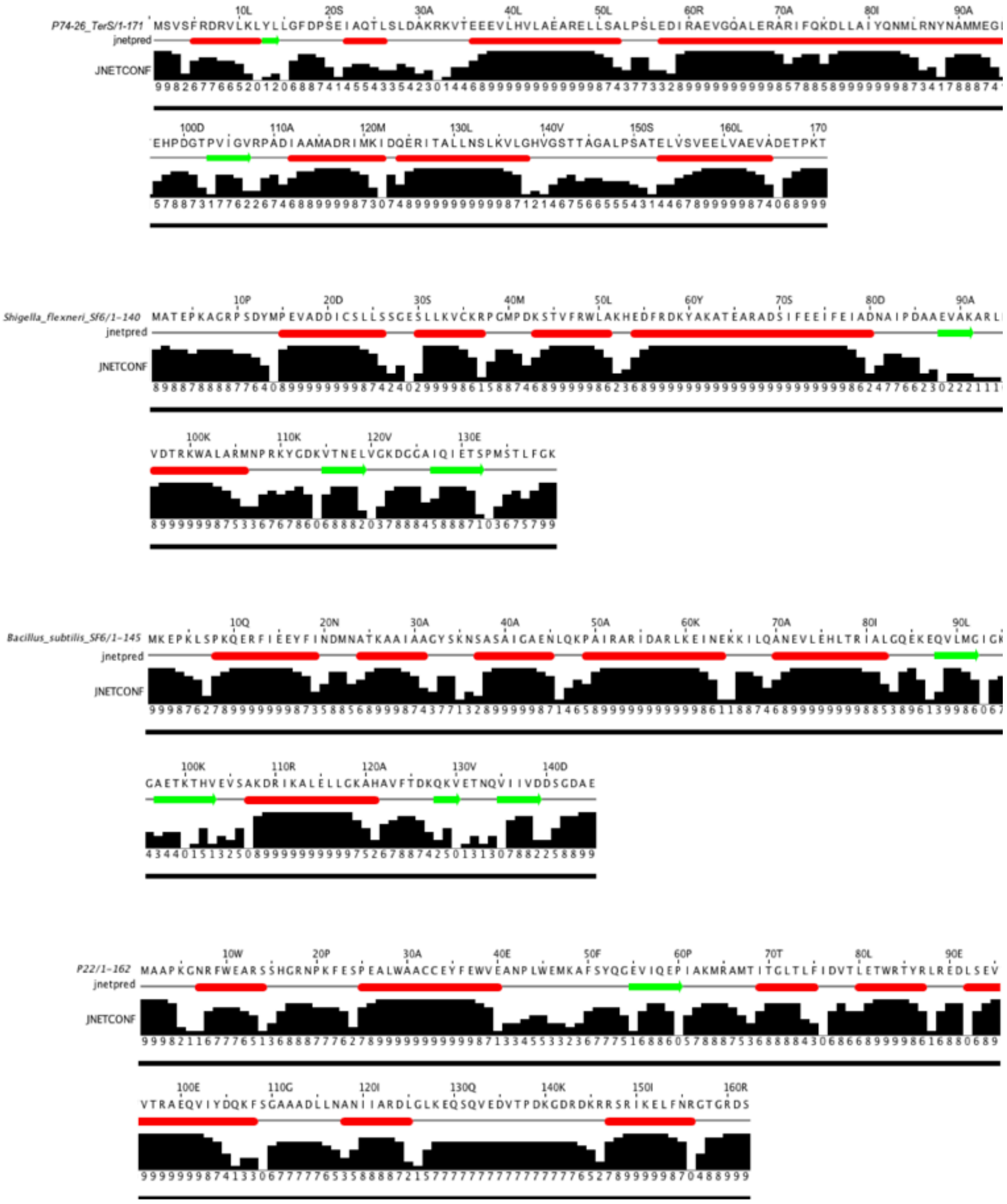

**Supplemental Figure 5. Secondary Structure Prediction of TerS$^{P74\text{-}26}$ .** Secondary structure was predicted using the JPred4 algorithm (Drozdetskiy et al 2015). 'jetnetpred' indicates the consensus prediction (red tubes for helical regions, green arrows for beta regions). 'JNETCONF' indicates the confidence estimate for the prediction (0-9). High values indicate high confidence, while low values indicate low confidence.

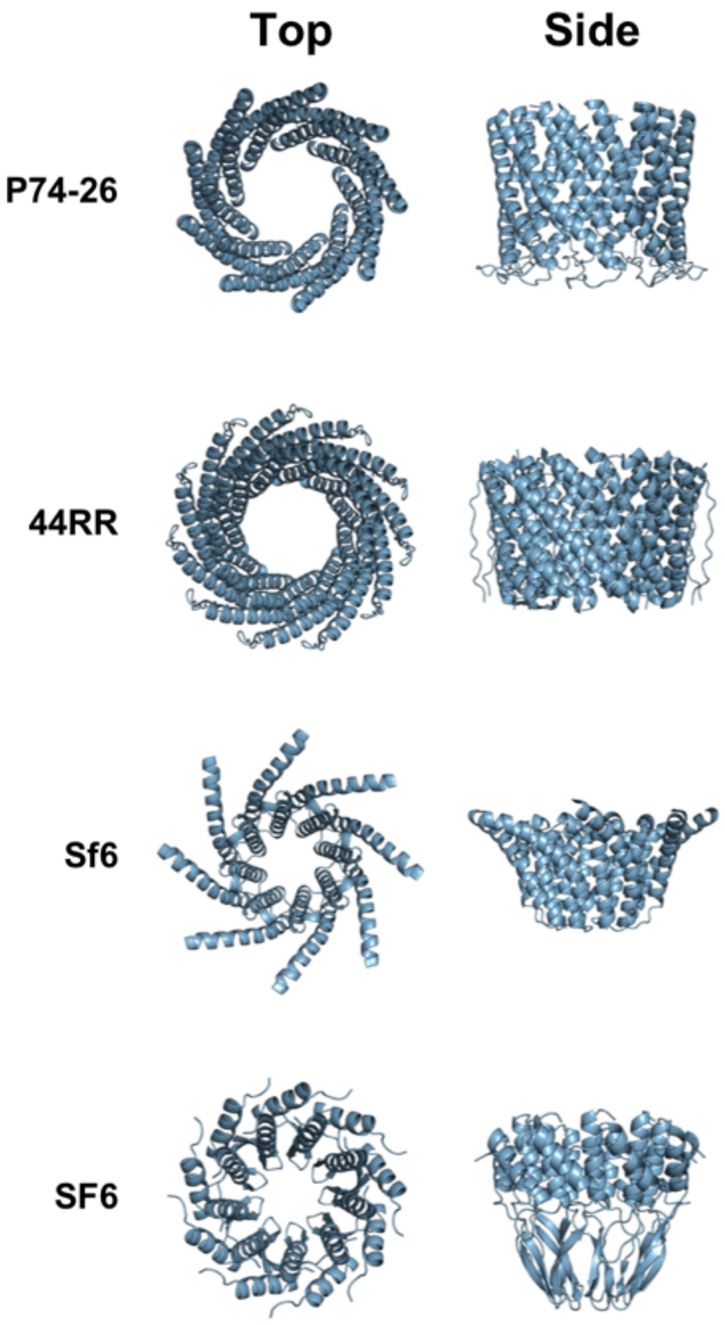


**Supplemental Figure 6. Comparison of TerS oligomerization domains.** Oligomerization domains are shown from the view of the C-terminal region (left) and side (right) for $TerS^{P74\text{-}26}$, $TerS^{44RR}$ (3TXQ), $TerS^{Sf6}$ (3HEF) and $TerS^{SF6}$ (3ZQQ).

```
                   1        10        20          30        40         50
SF6_TerS        ...KLSPKQERFIEEYFINDMNATKAAIA..AGYSKNSASAIGAENLQ.KPAIRARIDA
Sf6_TerS        PSDYMPEVADDICSLLS.SGESLLKVCKR.....PGMPDKSTVFRWLAKHEDFRDKYAK
P74_26_TerS     ...MSVSFRDRVLKLYL.LGFDPSEIAQTLSLDAKRKVTEEEVLHVLAE...ARELLS.
44RR_TerS       ...........................................................

                                  60        70         80        90
SF6_TerS        RL.K.....KIL.......QANEVLEHLTR.IALGQEKEQVLMGIGKGAETKTHVE...
Sf6_TerS        ATEA............RADSIFEEIFEIADNA...........................
P74_26_TerS     ...ALPSL...EDIRAEVGQALERARIFQKDL...LAIYQ.............NMLRNY
44RR_TerS       ........NRKIDQDDDYELVRRNMHYQSQML...LDMAK.............IALENA

                                            100       110             120
SF6_TerS        ..........................VSAKDRIKALELLGKAHAVFTD......KQKVE
Sf6_TerS        ..................IPDAAEV...AKARLRVDTRKWALARMNPRKYGDKVTNELV
P74_26_TerS     NAMMEGLTEHPDGTPVIGV.RPADIAAMADRIMKIDQERITALLNSLKVL.........
44RR_TerS       KNAD.............SPRHVEVFAQLMGQMTTTNKEMLKMHKEMKDLA.........

                           130
SF6_TerS        ......TNQVIIVDDS
Sf6_TerS        GKDGGAIQIETS....
P74_26_TerS     ................
44RR_TerS       ................
```

**Supplemental Figure 7. Structure-based sequence alignments of TerS structures.** Alignments of SF6 (3ZQQ), Sf6 (3HEF), P74-26, and 44RR (3TXQ) TerS proteins. Alignments performed by PROMALS3D (Pei et al., 2008). Figure generated using ESPript 3 (Robert and Gouet, 2014). Red characters outlined in blue are considered highly similar (Similarity score value >0.7).

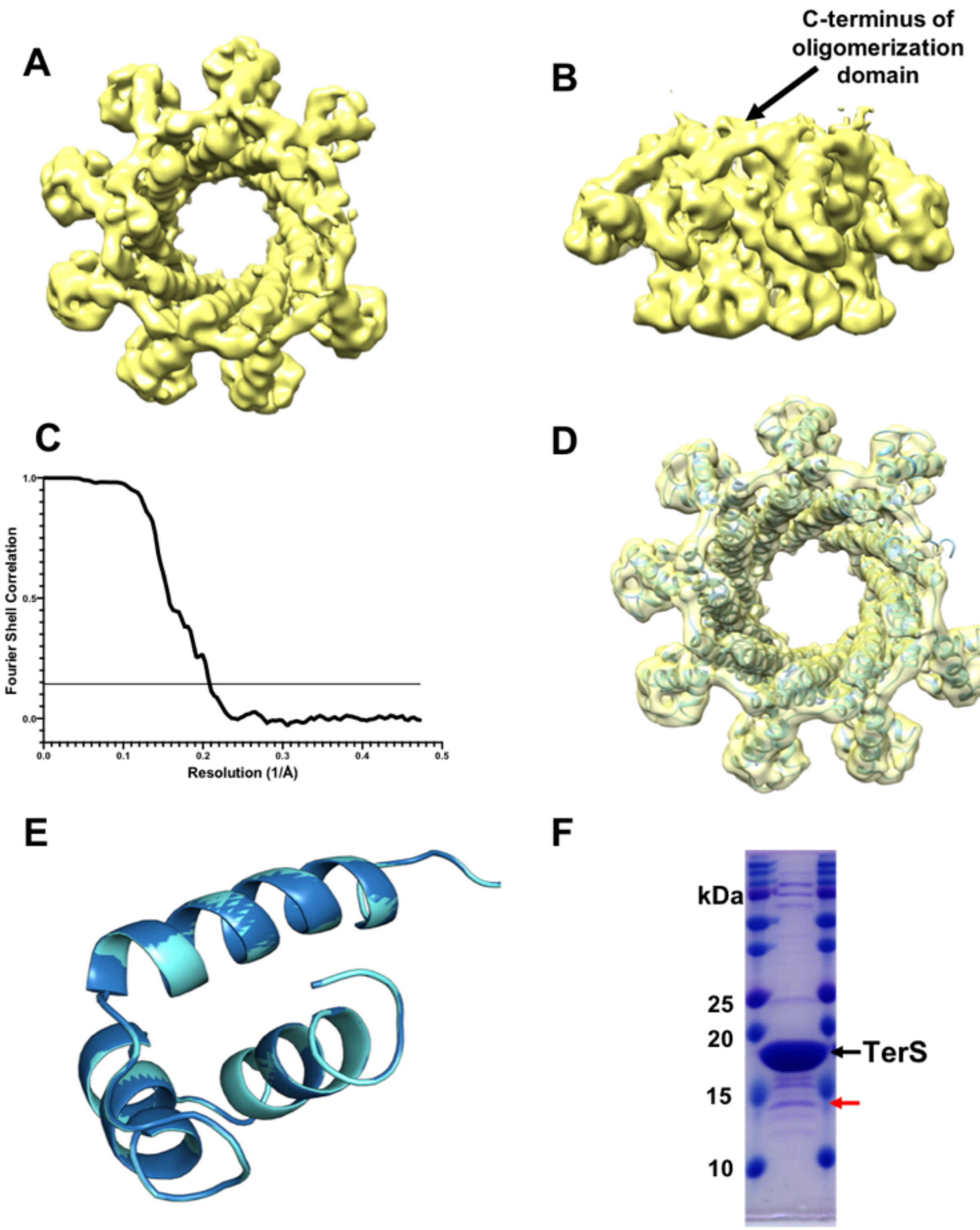


**Supplemental Figure 8. Asymmetric class 6 reconstruction of TerS$^{P74-26}$ . (A)** 4.8 Å resolution C1 asymmetric 3D reconstruction of the class 6 TerS ring (top) shows a missing HTH domain. **(B)** Side view of class 6 reconstruction. **(C)** Gold-standard FSC curve of class 6 3D reconstruction. Flat line represents the FSC 0.143 cutoff, indicating an overall resolution of 4.8 Å. **(D)** Class 6 atomic model in 3D reconstruction. **(E)** Superposition of chains A (dark blue) and C (light blue) of class 6 asymmetric model showing no conformational changes. **(F)** SDS-PAGE gel of P74-26 showing proteolysis (red arrow), the

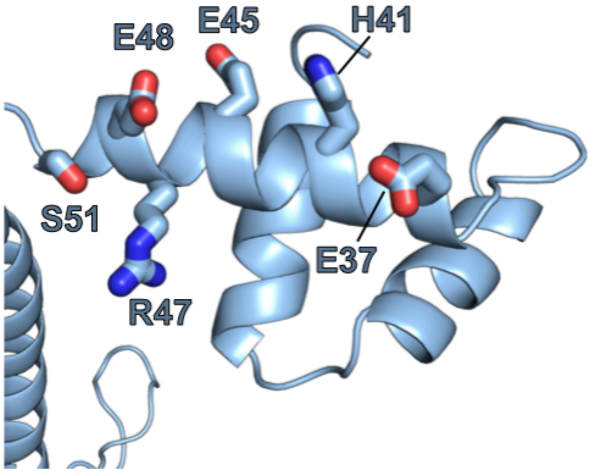


**Supplemental Figure 9. Charged and polar residues of helix 3.** Helix 3 is lined with several residues that can potentially bind the DNA major groove, including Glu37, His41, Glu45, Arg47, Glu48, and Ser51.

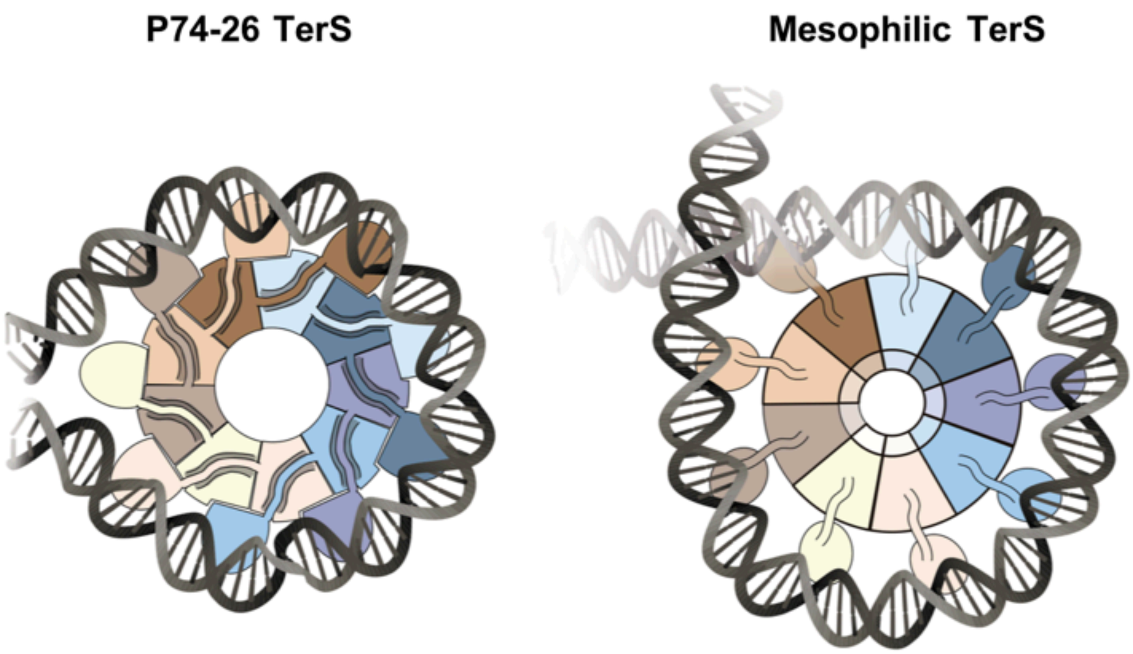


**Supplemental Figure 10. TerS$^{P74-26}$ and mesophilic TerS DNA wrapping models.** Rigid interactions between HTH domains and the barrel of the TerS$^{P74-26}$ ring forces the TerS$^{P74-26}$ complex to bind DNA in a planar conformation, disengaging one of the HTH subunits from DNA. In mesophilic TerS rings, the interactions between HTH domains and the barrel of the ring are weaker, allowing the HTH domains to adopt flexible conformations. These mesophilic TerS proteins may accommodate DNA superhelix formation, where the DNA spirals around the complex, staggering the HTH domains.

**Supplemental Table 1. Comparison of ringed TerS proteins with known structures.** PDB Codes: Sf6 (3HEF) SF6 (3ZQQ) P22 (3P9A) 44RR (3TXQ).

| Phage TerS | Pore diameter (Å) | HTH domain? | *Pac* length (nt) | C-terminus 2° structure | Favored DNA binding mode |
|---|---|---|---|---|---|
| P74-26 | 29 | Yes | ? | Predicted alpha | ? |
| Sf6 | 14 | Yes | 1800 | Beta | Wrapping |
| SF6/SPP1 | 18 | Yes | 242 | Beta | Wrapping |
| P22 | 20 | No | 22 | Beta/ predicted alpha | Threading |
| 44RR | 28 | ? | ? | ? | Wrapping |
